# Multi-omic gene regulatory networks informed by 3D chromatin architecture reveal insights into hematopoietic stem cell ageing

**DOI:** 10.64898/2026.04.19.719299

**Authors:** Quinn Stacpoole, Nadia Iannarella, Timothy M Johanson, Hannah D Coughlan, Rhys S Allan

**Affiliations:** The Walter and Eliza Hall Institute of Medical Research, Parkville, Victoria, 3052, Australia; Department of Medical Biology, The University of Melbourne, Parkville, Victoria, 3010, Australia

**Keywords:** 3D genome, haematopoietic stem cells, aging, epigenetics

## Abstract

During ageing, hematopoietic stem cells (HSCs) have reduced regenerative potential, skewed differentiation toward the myeloid lineage, and heightened susceptibility to clonal expansion and malignancy. While epigenetic alterations are well documented, the impact of aging on higher-order 3D chromatin architecture remains poorly understood. Here, we examined the 3D genome organisation of aged murine HSCs using in-situ Hi-C then integrated this with gene expression and chromatin accessibility data to build HiC-informed gene regulatory networks (GRNs). Aged HSCs display erosion of topologically associating domain (TAD) boundaries, A/B compartment switching, and reorganised enhancer-promoter loops associated with lineage-inappropriate gene expression. Our GRN analysis identifies a hierarchy of transcription factors, including a c-Maf-Lyl1-Mnt axis that orchestrates the transition from a youthful to aged state and a Gfi1-Sox4 axis in young HSCs that regulates Bach1. This study provides a structural blueprint for aging HSCs and defines specific regulatory targets for potential reprogramming interventions to restore hematopoietic youthfulness.

## Introduction

Hematopoietic stem cells (HSCs) are responsible for continually replenishing the blood system with mature cells, ensuring immune competence and tissue homeostasis (Osawa et al., 1996; Spangrude et al., 1988; Thomas et al., 1957). However, during the ageing process in mice and humans, HSCs expand due to increased self-renewal (Sun et al., 2014), decline in regenerative potential (Kuribayashi et al., 2021), shift toward myeloid-biased differentiation (Dykstra et al., 2011; Pang et al., 2011) and display increased genomic instability (Rossi et al., 2007; Rube et al., 2011), all of which contribute to reduced immune function and a higher risk of hematological malignancies in the elderly.

Studies have found that transplantation of aged murine HSCs into young mice can restore some function (Ergen et al., 2012; Montecino-Rodriguez et al., 2019; Yamamoto et al., 2018), especially after secondary transplantation (Yamamoto et al., 2018), but in many experiments transplantation of old HSCs into young mice does not correct the repopulating abilities of the stem cells (Dykstra et al., 2011; Liang et al., 2005; Rossi et al., 2005), suggesting an intrinsic dysfunction. HSCs accumulate epigenetic alterations with age, such as changes to chromatin accessibility (Itokawa et al., 2022) and DNA methylation (Bogeska et al., 2022) both of which alter gene expression in ways that reflect myeloid bias and a cellular reaction to an inflammatory bone marrow environment. However, whether these alterations in epigenetics and gene expression are primarily due to downstream effects of somatic mutations (Kapadia et al., 2025) or an independent reversable process of epigenetic alteration is still unclear since iPSCs derived from aged HSCs that are then redifferentiated to HSCs are functionally indistinguishable from young HSCs (Wahlestedt et al., 2017). Additionally, recent clonal tracing reveals that aged HSCs are likely an expanded population of epigenetically altered clones (Scherer et al., 2025).

A key driving factor in cell identity and gene regulation are the interactions between enhancers and promoters that are mediated via transcription factor (TF) binding (Martin et al., 2023), this is more broadly described as the 3 dimensional (3D) folding of the genome (Lafontaine et al., 2021; Tomas-Daza et al., 2023) and few studies have examined how the 3D genome of stem cells changes with age. Hi-C, a genome-wide chromatin conformation capture technique, has provided critical insights into the spatial organization of chromatin in mature immune cells (Bediaga et al., 2021; Chan et al., 2021; Johanson et al., 2018) and blood stem cells (Chen et al., 2019; Zhang et al., 2020). In addition to high resolution enhancer promoter interactions, Hi-C data allow for the identification of the large-scale hierarchical features of the 3D genome such as chromatin compartments (active A and inactive B compartments), topologically associating domains (TADs). Differential style analysis allows us to identity differences in the 3D genomes of conditions i.e. young and aged, and determine which regulatory elements and TFs are involved in the ageing of chromatin organisation (Lafontaine et al., 2021; Tomas-Daza et al., 2023). Studies in other cell types have revealed significant age-associated changes in chromatin organization. Age-related chromatin changes are often cell type-specific but share common features such as loss of TAD integrity, altered enhancer-promoter interactions, and redistribution of chromatin between A and B compartments (Kriukov et al., 2024; Ma et al., 2024). In muscle stem cells, aging is accompanied by a gain in long-range chromatin contacts and a loss of short-range interactions, impairing the transcription of regeneration-associated genes (B. A. Yang et al., 2023; Zhao et al., 2023). Neuronal aging, on the other hand, is characterized by the decay of heterochromatin domains, activation of transposable elements, and nuclear expansion (Kriukov et al., 2024; Zhang et al., 2022). Furthermore, Liu et al. demonstrated increased Shannon entropy in the B compartments of late-passage mesenchymal stromal cells (MSCs), indicating an increase in disorder with replicative age (Liu et al., 2022). Importantly, environmental and genetic interventions such as exercise and epigenetic reprogramming have been shown to mitigate some of these chromatin changes, emphasizing their potential reversibility (J.-H. Yang et al., 2023).

Often gene regulation is examined by correlating gene expression with the closest open chromatin region to infer gene regulatory elements. However, Hi-C allows us to directly link gene regulatory elements (often distal) to target genes (Javierre et al., 2016; Milevskiy et al., 2023). TF activity (i.e. the extent to which it is exerting its regulatory potential on its target genes through motif binding) has been shown to be a better predictor of cell-defining gene regulation than expression alone (Wlodarczyk et al., 2025). Even at low resolution, Hi-C data can add an additional layer during multiomic integration for mapping gene regulatory mechanisms and identifying TFs that govern gene regulatory networks (GRNs) (Fulco et al., 2019; Hecker et al., 2023). For example, integration of Hi-C, ATAC-seq and RNA-seq is one method of constructing GRNs, where TFs and genes are nodes and the strength of their interaction and activity are the edges (Fulco et al., 2019; Hecker et al., 2023). Furthermore, the differential analysis of network weights between conditions (e.g. cell type 1 vs cell type 2, or young vs aged) can provide additional information beyond differential expression or differential chromatin accessibility alone, since genes may change their connectivity through the 3D genome in the network with little change in their expression, allowing for the identification of TFs that drive network state changes (Hecker et al., 2023; Wlodarczyk et al., 2025; Xu et al., 2021). While the use of differential network analysis to identify TFs that drive trans-differentiation of one cell type to another is relatively common, its use to identify TFs that define aged states in the same cell type is a growing area of research (Lee et al., 2021; Randhawa & Kumar, 2021; Wang et al., 2023). However, none have yet used Hi-C derived genome organisation to inform the construction of TF-gene networks.

Investigations into the chromatin architecture of HSCs, let alone aged HSCs, are scarce, but in blood progenitors such as aged pro-B cells, alterations in enhancer-promoter interactions and shifts in chromatin compartmentalization have been linked to impaired B cell development (Ma et al., 2024). These findings imply that changes in chromatin architecture could play a role in HSC aging, yet the specific changes occurring in HSCs remain to be elucidated. Here, we examine the 3D genome organisation of young and aged murine long-term (LT-) HSCs using Hi-C to gain further insights into the molecular drivers of their functional decline and potentially identify novel strategies to reprogram aged HSCs back to a more youthful state. We find that ageing appears to cause a loss of TAD boundary strength and an increase in disorder in the 3D chromatin contacts of murine LT-HSCs. Additionally, differentially expressed genes (DEGs) are enriched in the differentially interacting (DI) regions (i.e. regions that gain or lose chromatin interactions with age) and a small set of upregulated genes appear to be lineage inappropriate, as they are not normally expressed in any young blood progenitors, but are expressed in aged LT-HSCs and are particularly enriched in the DI regions. Furthermore, while differential TF-gene networks are often used to identify TFs for direct cell reprogramming (Xu et al., 2021), we used similar methods to identify TFs that govern the TF-gene regulatory network states of aged and young LT-HSCs and our analysis predicts a potential role for c-Maf, Cebpd and Atf3 as top drivers of the aged state, while Gfi1, Hif3a, Sox4 and Hes1 are top TFs predicted to govern the youthful state.

## Results

### Mitochondrial membrane potential status does not confer differences in 3D genome architecture, but age does

To tackle potential heterogeneity of the aged LT-HSC pool we leveraged knowledge from a previous study that demonstrated that mitochondrial membrane potential (MMPot) status could discriminate between young-like LT-HSCs in old mouse bone marrow (Mansell et al., 2021). To this end, aged and young LT-HSCs (CD34^-^ CD48^-^ Flt3^-^ CD150^+^ LSKs) were sorted based on MMPot status (low and high) (see Methods and Fig. 1A) and subjected to Hi-C. We processed the data and performed differential analysis using diffHic (A. T. Lun & G. K. Smyth, 2015) between old and young then high and low MMPot groups. While aged mice had a higher proportion of MMPot low LT-HSCs than young mice, as previously reported (Mansell et al., 2021) (Supplementary Fig. 1B-C), there were no significant differential interactions (DIs) between MMPot High and MMPot Low LT-HSCs from either young or aged mice (Supplementary Fig. 1D). However, when contrasted according to age, both MMPot high and low LT-HSCs groups each displayed differences in chromatin contacts (Supplementary Fig. 1D), suggesting ageing has a much stronger effect on chromatin structure than MMPot status. Therefore, all further analysis was performed on LT-HSCs in which MMPot high and low groups were pooled to compare LT-HSCs by age only.

**Figure 1.**
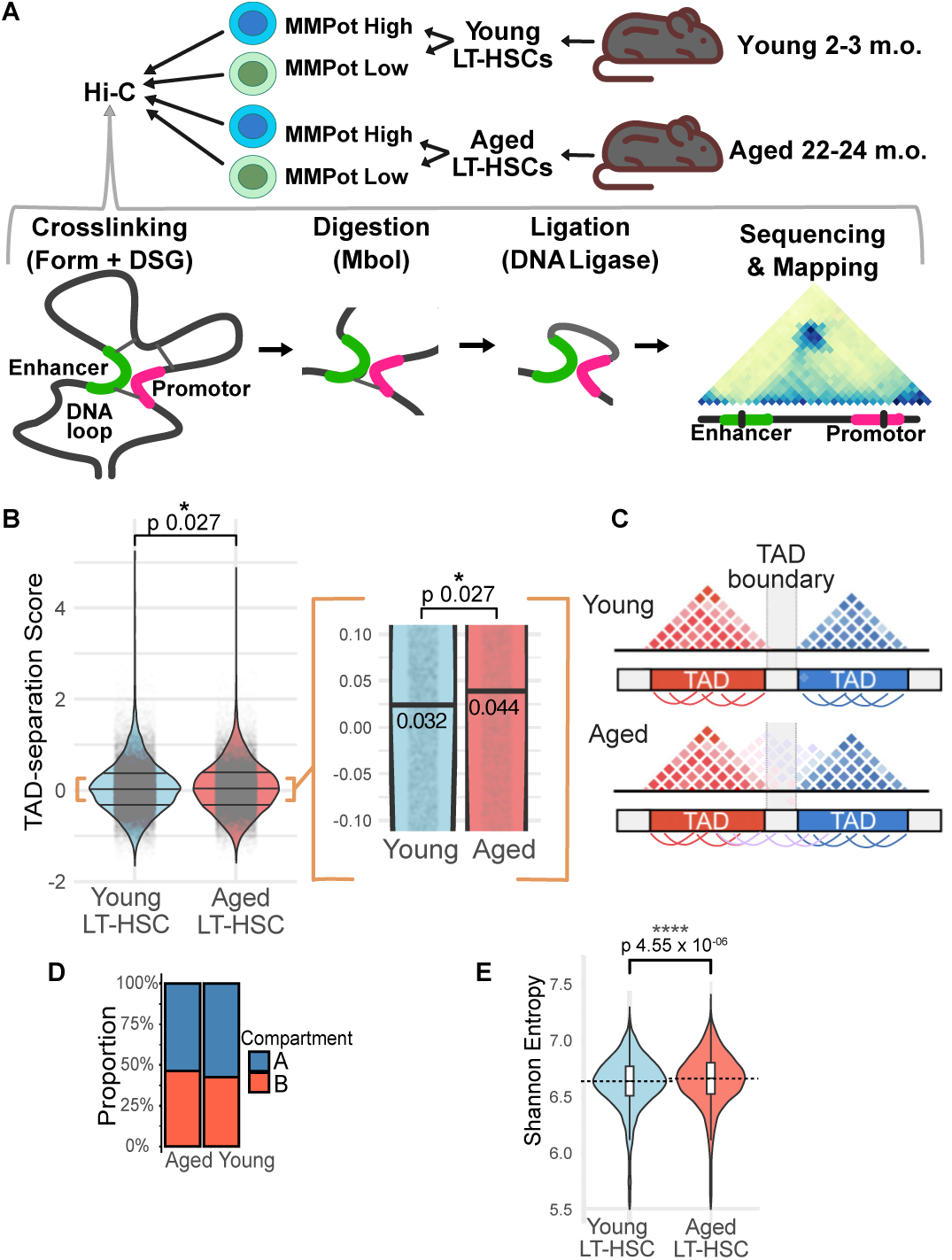
Age-associated changes in 3D chromatin architecture in LT-HSCs reveals increased disorder and a preponderance of gained interactions. A. Schematic of Hi-C experiment on LT-HSCs from young and aged C57Bl6 mice (MMPot = mitochondrial membrane potential, HSC = hematopoietic stem cell, Form = formaldehyde, DSG = disuccinimidyl glutarate); B. Violin plots of TAD separation scores in Young vs Aged LT-HSCs (difference assessed by Mann-Whitney U-test); C. Diagram depicting how an increase in TAD-separation score translates to overall reduced TAD boundary strength with more interactions occurring across TAD boundaries; D. A/B compartment proportions for Aged & Young LT-HSCs; E. Violin plots of Shannon entropy values for all 2 Mbp genomic spans in Young vs Aged LT-HSCs (difference assessed by Mann-Whitney U-test).

### Changes in higher order chromatin structures indicate an age-associated increase in disorder

To understand the higher order genomic structures, we assessed the number and strength of TADs using HiCExplorer (Ramirez et al., 2018) in which TAD boundary “separation scores” are calculated to call TADs. In our data, TAD separation scores are increased in aged LT-HSCs (Fig. 1B & C; Supplementary Table 1), indicating that there are more interactions across TAD boundaries, i.e. a loss of TAD boundary strength. Segments of the genome can also be classified into A or B compartments depending on how the large-scale interactions segregate within principal component analysis. Calculation of A/B compartments in our LT-HSCs using HiCDOC (Kurylo et al., 2023) reveals an age-associated increase in the proportion of B compartments and decrease in the proportion of A compartments (Fig. 1D; Supplementary Table 1).

We wondered if the age-associated loss of TAD boundary strength reflects a general increase in disorder in the LT-HSC chromatin structure. To address this, we calculated Shannon entropy for young and aged LT-HSCs by calculating the probability distribution of contact frequencies for every 2 Mbp genomic span tiled along the mm39 genome, a method previously used in other models of ageing (Jenkinson et al., 2017; Liu et al., 2022). Aged LT-HSCs have a higher median Shannon entropy across all 2Mbp spans than young LT-HSCs (Fig. 1E; Supplementary Table 1), indicating a subtle erosion of the contact frequency landscape with ageing. Altogether, the changes in TAD boundary strength, changes to compartments and the overall increase in Shannon entropy suggest an increase in the disorder of the chromatin architecture in aged LT-HSCs.

### Aged LT-HSCs develop chromatin interactions associated with differentially expressed genes

DI analysis was performed on aged versus young LT-HSCs using, which revealed that 1353 cis-interactions decrease in interactivity and 3448 increase in interactivity during ageing in murine LT-HSCs (Fig. 2B, Supplementary Table 2). The highest ranked region (by false discovery rate (FDR)) in the analysis, was a gain in interactions on chromosome 5 between a 100 kbp region near *Gabra2* and a 100 kbp region upstream near *Guf1* (Fig. 2A & C, left panel). Interestingly, many chromosomes 5 regions near other GABA receptor ion-channel components *Gabrb1* and *Gabra4* also gained interactions (Fig 2G, middle panel). The highest ranked lost interaction was on chromosome 15 near *Nipbl* and *Wdr70*, respectively (Fig. 2C, right panel). To compliment this analysis, we reanalysed previously published RNA-seq data from Itokawa et al 2022 data (see Methods) to determine differentially expressed genes (DEGs) between young and old HSCs. We observed that DEGs are significantly enriched in the DI regions (Fig. 2E). Furthermore, the DEGs *B3galt1*, *Gabra4*, *Gabra1* and *Terf1* all overlap with the top 20 DI regions that alter interaction frequency with age (Fig. 2G and Supplementary Fig. 2A). Interestingly, comparison of aged versus young murine monocytes or natural killer (NK) cells showed a negligible number of DIs at 100 kbp resolution (Supplementary Fig. 1F), suggesting that either mature blood cells correct their 3D genome as they differentiate, or that the myeloid and lymphoid progenitors that go on to differentiate to maturity already have chromatin architecture similar to that of young progenitors.

**Figure 2.**
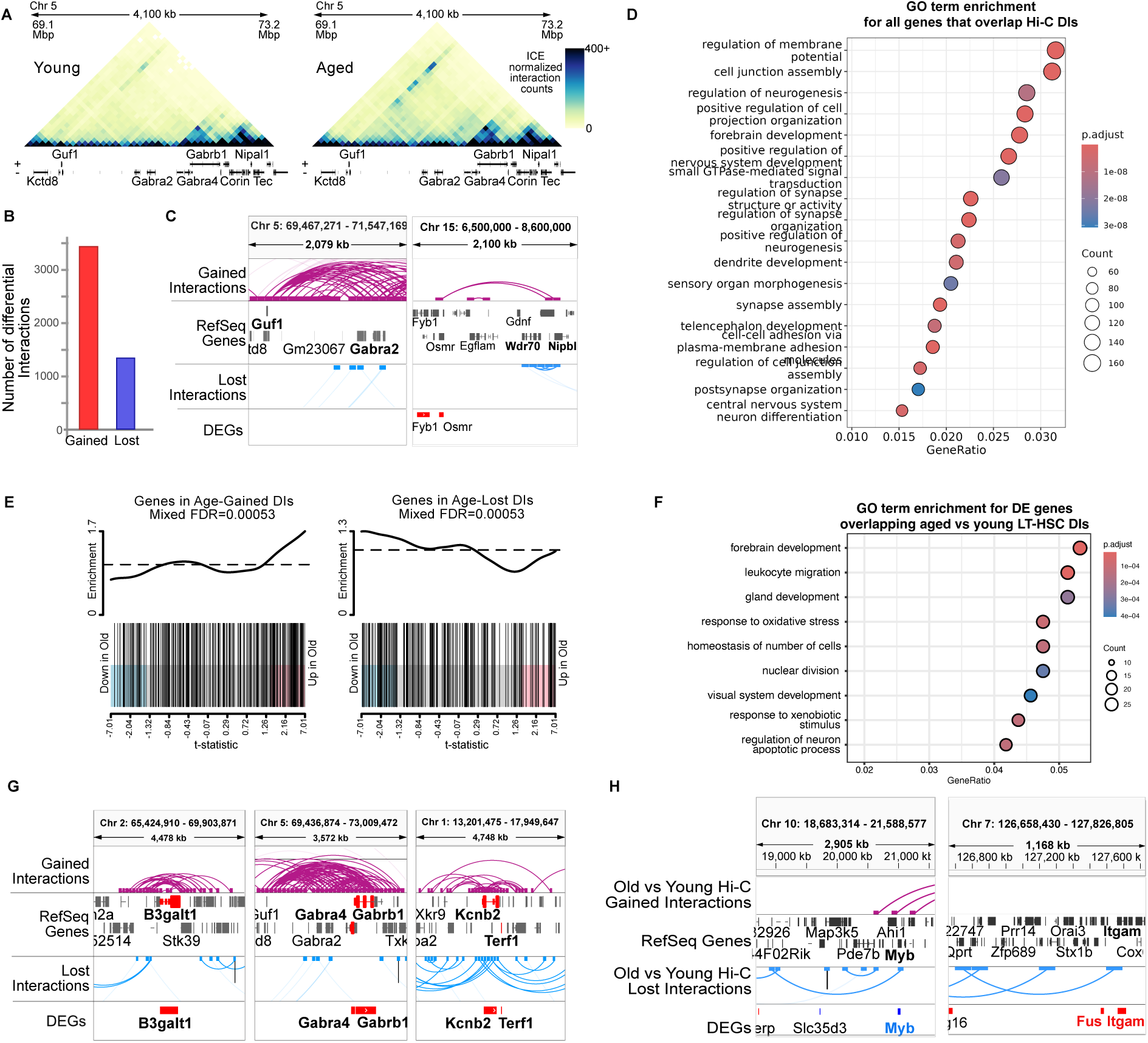
Genes overlapping regions which alter their interactions with age in murine LT-HSCs reveals biological processes associated with signalling, activation, migration and differentiation. A. Hi-C Contact maps (bin size = 100 kbp) of the region 69.1 – 73.2 Mbp on chromosome 5 for Young and Aged LT-HSCs, which includes the most significant differential interaction (DI) that occurs between regions around Guf1 and Gabra2; B. The number of differential intra-chromosomal (cis) interactions in Aged/Young LT-HSCs (red bar = interactions gained with age, blue bar = interactions lost with age); C. Arc plots of the most significant (FDR < 0.1) gained (left panel) and lost (right panel) DIs at 100 kbp resolution; D. GO term enrichment for all genes that overlap Hi-C aged vs young LT-HSC DIs; E. Enrichment of differentially expressed genes (DEGs) in aged vs young DIs using the fry test where “mixed” false discovery rate (FDR) is the FDR value for enrichment in the combined gained and lost DIs; F. GO term enrichment for DEGs overlapping aged vs young LT-HSC DIs; G. Arc plots of the most significant gained and lost DIs at 100 kbp resolution and the upregulated DEGs they overlap (upregulated DEGs shown in red). H. Age-associated loss of interactions between a region near myeloid-associated gene *Myb* and a region ∼2 Mbp downstream of Myb’s promotor (1st panel) and age-associated loss of interactions between a region overlapping myeloid-associated gene *Itgam* and a region over 800 kbp upstream (2nd panel).

Gene Ontology (GO) term enrichment analysis of genes near regions that change chromatin architecture with age in LT-HSCs revealed an enrichment of genes involved in the regulation of membrane potential, cell junction assembly and many genes involved in neurogenesis (Fig. 2D; Supplementary Table 3), likely reflecting share processes. Kyoto Encyclopedia of Genes and Genomes (KEGG) pathway enrichment analysis revealed many genes involved in cancer (especially miRNAs), as well as glutamatergic and serotonergic synapses (Supplementary Fig. 2B; Supplementary Table 3). When GO term analysis is restricted to only genes that are both differentially expressed (from the Itokawa et al 2022 dataset) and that also overlap with regions of differential interactions, many genes appear to be involved in leukocyte migration (especially myeloid cell migration) and neuron-related processes (Fig. 2F; Supplementary Table 3), while efferocytosis pathway genes were the most significantly enriched in Kyoto Encyclopedia of Genes and Genomes (KEGG) analysis (Supplementary Fig. 2C; Supplementary Table 3). As expected, many prominent genes involved in LT-HSC activation and myeloid differentiation appear in the GO term enrichment and show both age-associated chromatin contact changes and changes in gene expression (Supplementary Table 3). For example, *Myb* (aka *c-Myb*) and *Itgam* (among others) appears in GO terms such as “myeloid cell homeostasis”, “myeloid leukocyte migration” and “negative regulation of myeloid cell differentiation” to name a few (Supplementary Table 3). The expression of *Myb* decreases with age in LT-HSCs as does the frequency of interactions between a region near *Myb* and a region ∼2 Mbp upstream of *Myb*’s promotor (Fig. 2H, 1^st^ panel). Similarly, *Itgam* (*Cd11b*) is upregulated in aged LT-HSCs and a region overlapping its promotor shows a decreased frequency of interactions with a region over 800 kbp upstream (Fig. 2H, 2^nd^ panel).

### Genes that have age-related inappropriate lineage expression overlap with changes in genome organisation

We reasoned that the observed increase in disorder within the chromatin contacts might result in the expression of genes that are not usually expressed in the blood-cell progenitor lineage. To test this, we used the raw counts from the Itokawa et al. (2022) RNA-seq data, which contains expression data for young (8 week old) and aged (20 month old) blood progenitors, including LT-HSCs, multipotent progenitors 1-4 (MPP1-4s), megakaryocyte-erythroid progenitors (MEPs), granulocyte-monocyte progenitors (GMPs) and common lymphoid progenitors (CLPs). We defined a set of genes that have low or negligible expression in young LT-HSCs, MPP1-4s, GMPs, MEPs or CLP and significantly higher expression in aged LT-HSCs (Supplementary Fig. 3B), which resulted in a list of 85 genes (Supplementary Fig. 3A). GO term enrichment of these DEGs that are very lowly or absent in young progenitors highlighted groups of genes such as *Tmem215*, *Kdr*, *Bmp4*, *Myo6*, *Ror2*, *Tbx1* and *Rorb* being involved in sensory organ morphogenesis and *Kdr, Bmpr1a, Bmp4 and Tbx1* involved regulation of mesenchymal cell proliferation (Supplementary Fig. 3D and E). Restricting our definition again to genes that have very low expression in young blood progenitors (Supplementary Fig. 3C) defined a list of genes we term “lineage inappropriate genes” (LIGs) which refined our list to 8 genes: *Rcvrn*, *Ntf3*, *Syt15*, *Mab21l2*, *Rorb*, *Aldh3a1*, *Chrm3* and *Tdrd9* (Fig. 3A & C). Interestingly, 7 out of 8 strict LIGs (which are also DEGs) also overlap with and are significantly enriched within regions of differential interactions in our Hi-C data compared to other randomly selected sets of upregulated DEGs (Fig. 3B), likely indicating that age-related 3D chromatin changes lead to expression of these lineage inappropriate genes.

**Figure 3.**
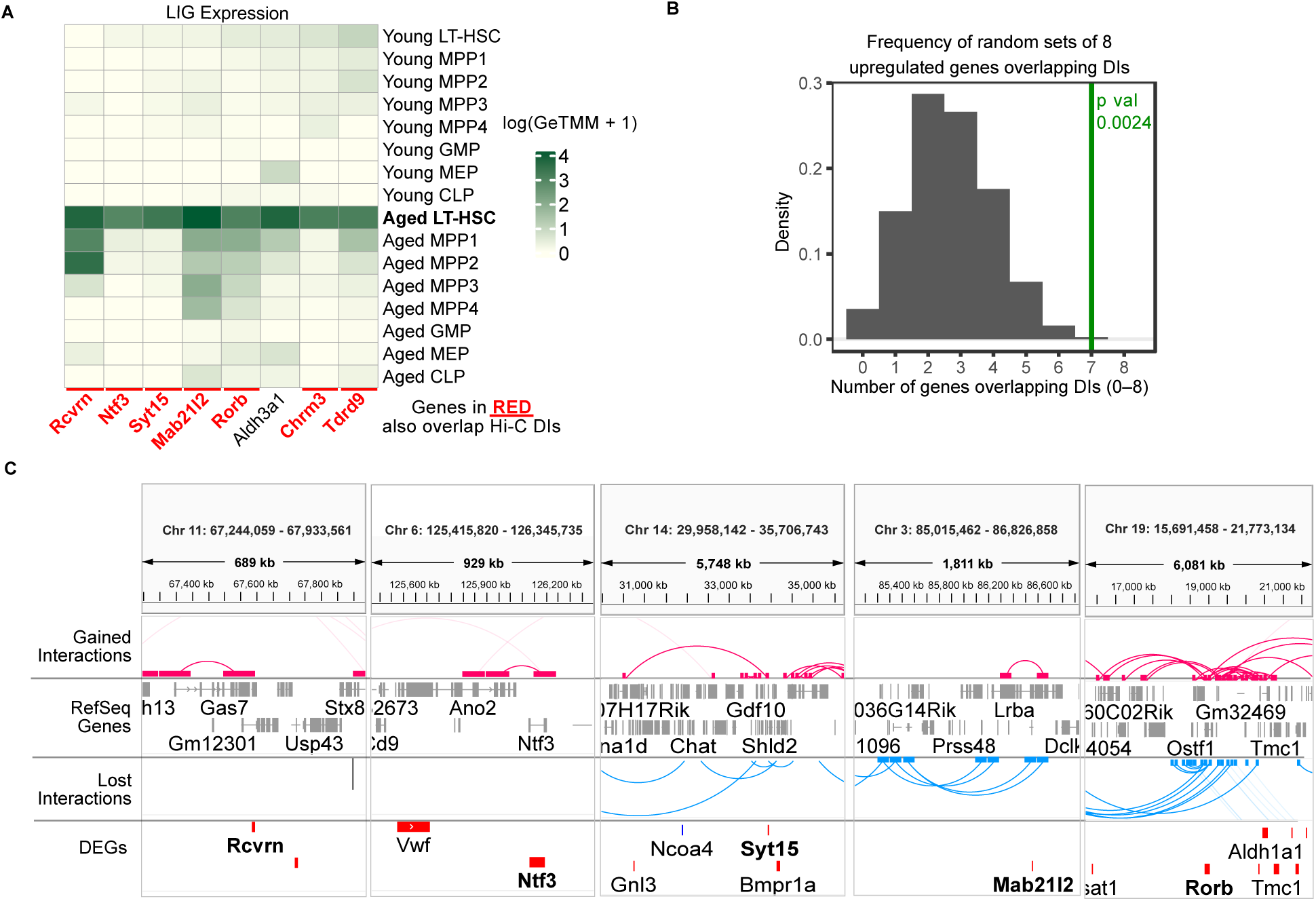
Genes not normally expressed in any young murine blood progenitors are upregulated with age, especially in LT-HSCs, and 7/8 overlap aged vs young LT-HSC Hi-C DIs (genes in red). A. Heatmap of expression (logGeTMM+1) across all blood progenitors for the 8 strictly defined lineage inappropriate genes (LIGs) identified in the Itokawa et al 2022 dataset (genes in green and columns marked with a green bar also overlap aged vs young LT-HSC differential interactions (DIs); B. Density plot of how often genes within sets of 8 randomly chosen upregulated genes (testing 10000 sets of 8 genes) overlap DI regions as determined by a random sampling Monte Carlo permutation test (the p value of the likelihood of 7 upregulated genes overlapping DIs is 0.0024 as determined by the Monte Carlo empirical test); C. Interaction arc plots for gained and lost interactions for aged vs young LT-HSC DIs (100 kbp resolution) that overlap 5 of the LIGs: *Rcvrn*, *Ntf3*, *Syt15*, *Mab21l2* and *Rorb*.

### AP-1 factor binding motifs are enriched in chromatin that opens and gain interactions during ageing

We wondered if regions that change in interaction frequency are enriched for particular TF motifs. To answer this question, we used ATAC-seq data to determine differentially accessible regions (DARs) between young and aged LT-HSCs using the Itokawa et al 2022 data, then performed motif enrichment analysis within DARs that overlap our Hi-C DI regions (Fig 4A). Strikingly, motifs for AP-1 factors Jun, Jund and Fosl2, as well as the common AP-1 binding partner Ets1, are enriched in age-opening peaks that overlap regions that gain interactions (Fig. 4B, age-opening/age-gained columns). There is also a mild enrichment of PPARG and NR1H3 motifs in age-closing peaks that overlap regions that lose interactions with age (Fig. 4B, age-closing columns).

**Figure 4.**
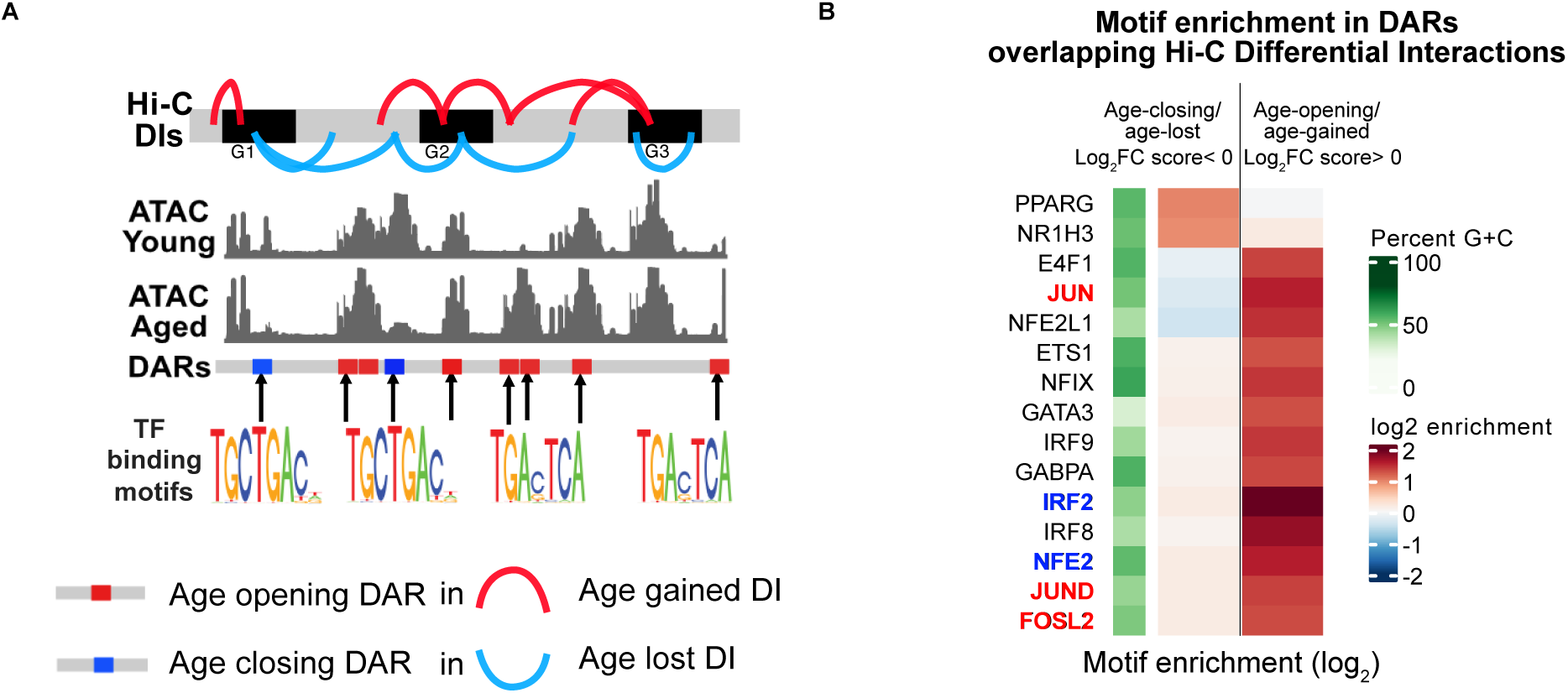
Differentially accessible regions (DARs) overlapping DIs with the same logFC direction show strong enrichment for AP-1 factor motifs. A. Schematic of motif enrichment analysis whereby transcription factor (TF) binding motif enrichment was assessed in age-opening and age-closing DARs that overlap Hi-C differential interactions with the same log fold change sign (e.g. logFC > 0 means age-opening DARs in gained DIs, logFC < 0 means age-closing DARs in lost DIs); B. MonaLisa TF motif enrichment results showing the top HOCOMOCO v13 motifs enriched in age-closing DARs that overlap age-lost differential interactions (Log_2_FC score < 0) and in age-opening DARs that overlap age-gained differential interactions (Log_2_FC score > 0) – red highlighted TFs are upregulated in aged LT-HSCs, blue highlighted TFs are downregulated.

### Integrated multiomic analysis predicts TFs that drive the aged and youthful states of LT-HSCs

To further explore how transcription factor binding influences age-related changes in epigenetics and gene regulation we used differential GRN analysis. While our motif enrichment analysis highlighted the DEGs Jun, Jund and Fosl2, TFs regulate genes in a network and can change network connectivity with little change in expression (Wlodarczyk et al., 2025). As such, changes in network connectivity can reveal TF drivers of network state change. Moreover, analysis of differential GRNs can rank TFs based on the strength of the change in connectivity to predict which TFs to use for cell reprogramming (Kamaraj et al., 2020; Rackham et al., 2016; Xu et al., 2021). Therefore, we constructed GRNs using our Hi-C data and publicly available ATAC-seq and RNA-seq data using a combined workflow of STARE (Hecker et al., 2023) and a modified pipeline similar to ANANSE (Xu et al., 2021) (Fig 5A).

**Figure 5.**
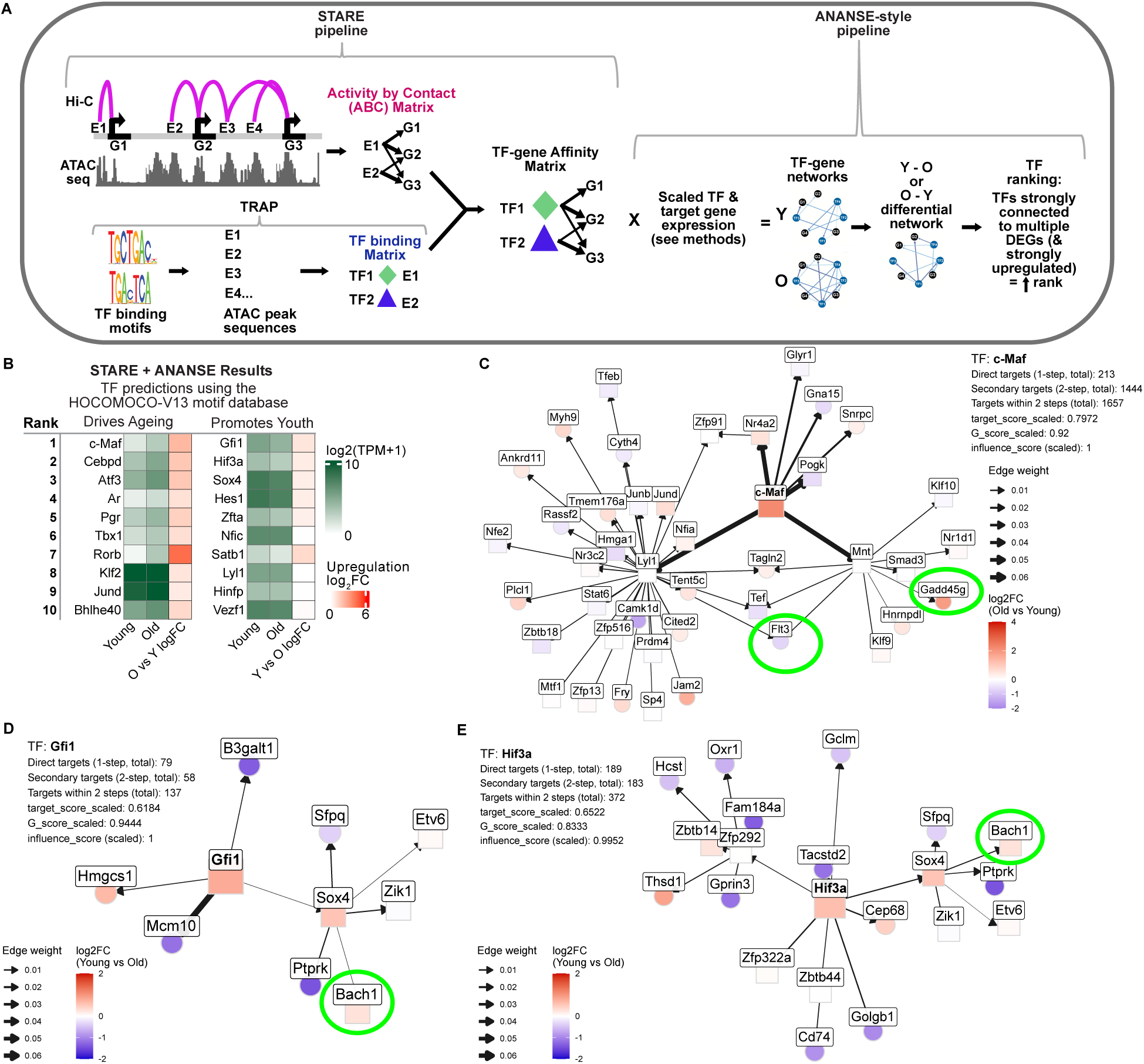
Integrated differential network analysis predicts transcription factors (TFs) that drive the aged and young states of murine LT-HSCs. A. Schematic of the combined STARE and ANANSE-style pipeline that integrates Hi-C, ATAC-seq and RNA-seq data: purple arcs represent ICE-normalized Hi-C interaction frequencies that overlap candidate enhancers (E1-E4), and genes (G1-G3) 2.5 kbp either side of the gene’s TSS, which are combined with ATAC-seq loess offset normalized peak values to generate an activity by contact (ABC) matrix, which is combined with a TF binding matrix to generate a TF-gene affinity matrix, which is then used by the ANANSE-style pipeline to generate differential gene regulatory networks (GRNs) and rank the TFs that drive the GRN changes; B. TF ranking results using the HOCOMOCO v13 TF motif database showing the expression (log2 TPM+1) and fold-change (log2FC) heatmaps of the top 10 ranked TFs predicted to either (i) drive the gene regulatory network towards the aged state (left 3 “Drive Ageing” columns) or (ii) drive the network towards the young state (right 3 “Promotes Youth” columns); C. Simplified network diagram of the top ranked age driving TF, c-Maf, which is predicted to regulate 213 direct targets in this GRN; D. Simplified network diagram of the top ranked youth promoting TF, Gfi1, which is predicted to regulate 79 direct targets, including Sox4; E. Simplified network diagram of the 2^nd^ top ranked youth promoting TF, Hif3a, which is predicted to regulate 189 direct targets, including Sox4

The top 10 ranked TFs predicted to drive each condition are shown in Fig. 5B and Supplementary Table 4; c-Maf (aka Maf), Cebpd, Atf3, Ar, Pgr, Tbx1, Rorb, Klf2, Jund and Bhlhe40 are predicted to drive the gene regulatory network changes during LT-HSC ageing, while Gfi1, Hif3a, Sox4, Hes1, Zfta, Nfic, Satb1, Lyl1, Hinfp and Vesf1 are predicted to be candidates to restore the youthful LT-HSC gene regulatory network state.

Inspection of these Hi-C informed differential GRNs (Supplementary Table 4) predicts that c-Maf regulates the TFs Lyl1 and Mnt to influence Flt3 downregulation in aged LT-HSCs (Fig. 5 C) as does Cebpd via Myc (Supplementary Fig. 4A). Both c-Maf1 via Mnt (Fig. 5C) and Cebpd via Myc (Supplementary Fig. 4A) are also predicted to drive the upregulation of Gadd45g, a driver of differentiation (Thalheimer et al., 2014). Furthermore, both Cebpd and Atf3 are predicted in our differential GRN to directly downregulate the TF Bach1 (Supplementary Figs. 4A and B), a known suppressor of myelopoiesis (Itoh-Nakadai et al., 2014; Kato et al., 2018). The LIG Rorb is ranked 7^th^ as a driver of LT-HSC ageing (Fig. 5B) and is predicted in the ageing-specific differential GRN to drive the downregulation of the TFs Dbp & Nr4a1 (Supplementary Fig. 4C).

Inspection of the Hi-C informed differential GRNs driving the youthful LT-HSC state predicts that Gfi1 & Hif3a both target Sox4 to upregulate Bach1 (Fig 5 D & E; Supplementary Fig. 4D), previously mentioned as a suppressor of myelopoiesis, and therefore Gfi1 & Hif3a have the opposite role of Cebpd and Atf3, which were predicted to downregulate Bach1. The 4^th^ ranked TF predicted to drive the young LT-HSC GRN is Hes1 (Fig. 5B) and inspection of its immediate targets shows it is predicted to suppress the expression of the sphingosine-1-phosphate transporter gene, *Mfsd2b*, and upregulate the suppressor of cholesterol synthesis gene, *Insig1* (Supplementary Fig. 4E).

It is expected that many DEGs in the networks would overlap our Hi-C DIs and inspection of the target DEGs of the top ranked age-driving TF, c-Maf (Fig. 5A), show that many do, including the DEGs Glyr, Plcl1, Tent5c, Zbtb18, Jam2 and Fry (Supplementary Fig. 4F).

## Discussion

Our results are, to our knowledge, the first to show that the 3D chromatin architecture of aged murine LT-HSCs is altered and demonstrates that aged LT-HSCs display a loss, albeit subtle, of overall TAD boundary strength and an increase in chromatin interaction Shannon entropy, which indicates that there is a general increase in disorder within the 3D chromatin structures. This increase in chromatin interaction Shannon entropy is in line with *in vitro* studies looking at late passage mesenchymal stem cells which show an increase in Shannon entropy in B compartments (Liu et al., 2022). We found negligible differences in chromatin architecture in aged vs young monocytes and NK cells. Since myeloid-biased LT-HSCs with low differentiation capacity are clonally expanded from a small set of original stem cells (Scherer et al., 2025), we hypothesise that LT-HSC fate is epigenetically set early in the mouse lifespan and that the aged monocytes and NK cells mostly derive from the remaining young-like LT-HSCs with youthful genome architecture.

Obtaining sufficient numbers of LT-HSCs is difficult and a limitation of our Hi-C data is the low resolution due to low cell input. Although we did not find any differences in chromatin architecture between LT-HSCs with high and low mitochondrial membrane potential, this does not exclude this possibility. Increased cell input, and thus higher resolution, may reveal more subtle differences in chromatin architecture between MMPot high and low LT-HSCs and other LT-HSC sub-populations such as CD150 high and low LT-HSCs which have been shown to be functionally distinct (Wang et al., 2025). Despite these caveats, our analysis was able to detect clear age-related 3D genome changes suggesting that MPPot status does not correlate with overt epigenetic alterations.

Since aged phenotypic LT-HSCs (CD34^-^ CD48^-^ Flt3^-^ CD150^+^ LSKs) contain a higher proportion of myeloid-primed LT-HSCs, locus-specific differences in chromatin are likely to reflect this myeloid priming. For example, our results show a region near the gene *Nipbl* has a strong loss of interactions with a region near *Wdr70* and a gain in interactions with a region near *Osmr* (Fig. 2C). *Nipbl* encodes a protein, Nipped B-like, a cohesin complex chromatin loading protein that is more abundant in aged LT-HSCs (Zaro et al., 2020) and likely upregulates *Runx1* and increases myeloid cell differentiation (Mazzola et al., 2020), thus supporting the observed increase in myeloid-biased LT-HSCs. *Osmr*, encodes a subunit of the oncostatin M (OSM) receptor and that is significantly upregulated in aged murine HSCs (Flohr Svendsen et al., 2021; Itokawa et al., 2022) and promotes inflammatory signalling (Jakob et al., 2021) and HSC mobilization (Bisht et al., 2022). The observed downregulation of *Myb* and upregulation of *Itgam* (*Cd11b*) are also known to be important for myeloid differentiation and these DEGs overlap differentially interacting regions in our aged vs young LT-HSCs (Fig. 2H). *Myb* is likely a target of the Runx1 transcription factor during differentiation of megakaryocyte-biased LT-HSCs (C. Wang et al., 2023) and its downregulation is important for differentiation of LT-HSCs into downstream lineages (Bianchi et al., 2010; Garcia et al., 2009; Sakamoto et al., 2012). *Itgam* upregulation in LT-HSCs is associated with both enhanced regenerative potential and myeloid cell differentiation (Rydstrom et al., 2023; Trento et al., 2017).

It is also interesting to note that regions near GABA receptor ion-channel genes *Gabra2*, *Gabrb1* and *Gabra4* show many gained interactions and *Gabrb1* and *Gabra4* are both upregulated. In addition, the top two terms in our DI GO term enrichment analysis were “regulation of membrane potential” and “cell junction assembly” and there were many other gene sets associated with ion transport and neurogenesis (Fig. 2 and Supplementary Fig. 2). A recent proposed model of ageing proposes ageing as a loss of the collective goal of cells mediated by changes in the bioelectric network, which is governed primarily by regulation of ion-channels (Pio-Lopez & Levin, 2024). A promising avenue of further research would be to determine if there is a difference in bioelectric patterns in young vs old bone marrow and whether the manipulation of ion-channels could restore old bioelectrical patterns in the bone-marrow to a younger state.

We determined so-called lineage inappropriate genes (LIGs) which are not expressed in young blood lineages but are upregulated in aged LT-HSCs, genes which are normally expressed in other lineages. For example, *Rcvrn* (coding for Recoverin) is normally expressed in retinal cells and yet is expressed in aged LT-HSCs but not in any young blood progenitors (Fig. 3A and 3C panel 1). Our finding that LIGs (which are also DEGs) show a disproportionate overlap with DI regions in the Hi-C data compared to other upregulated DEGs, suggests that the regions of chromatin around the LIGs are susceptible to age-associated 3D chromatin changes that lead to expression of these genes. We speculate that the expression of lineage inappropriate genes is a consequence of an overall increase in disorder of the chromatin interactions (TF mediated low-frequency enhancer-promoter interactions), which could allow for the selection of LT-HSCs in which these changes are advantageous for survival and proliferation. Examination of the mutational burden of aged murine LT-HSCs would be required to assess whether mutations linked to clonal hematopoiesis are also involved.

We also report a strong enrichment of AP-1 binding motifs in the DARs that overlap regions of age-gained interactions, in line with a recent paper demonstrating in 22 different murine cell types that regions that increase in chromatin accessibility during ageing show enrichment for AP-1 binding motifs (Patrick et al., 2024). Our novel multiomic integration pipeline uniquely integrates Hi-C interaction data with ATAC-seq and RNA-seq using a hybrid workflow of STARE (Hecker et al., 2023) and the TF prediction tool ANANSE (Xu et al., 2021). ANASE is normally used to predict TFs for direct cell reprogramming of one cell type to another, but which we have adapted to predict TFs that govern different states of LT-HSCs ageing. Wahlestedt and colleagues demonstrated that aged myeloid-biased LT-HSCs can be epigenetically reset to have a balanced output via reprogramming to a pluripotent state (Wahlestedt et al., 2017). However, there is a growing use of direct reprogramming factors to reinforce the cell identity lost with age (Roux et al., 2022) or other factors that promote youthful gene expression (Sengstack et al., 2026) or epigenetic patterns (de Lima Camillo et al., 2025) without any transition through a pluripotent state. From our Hi-C informed GRN analysis in LT-HSCs, the TFs most strongly predicted to drive the aged state are c-Maf, Cebpd & Atf3 and thus would be candidates for inhibition. On the other hand, Gfi1, Hif3a, Sox4 and Hes1 overexpression are predicted to be candidates to restore the youthful LT-HSC gene regulatory network state.

The results of our differential GRN analysis are strongly supported by the literature. For example, c-Maf family proteins have known roles in LT-HSC proliferation and maintenance (Asano et al., 2025; Hurt et al., 2004). As shown by us and others (Itokawa et al., 2022; Mooney et al., 2017), Flt3 mRNA expression is detectable in LT-HSCs (even though the protein is not present on the surface of during sorting) and is downregulated in aged LT-HSCs. Interestingly, our Hi-C informed differential GRN predicts that c-Maf regulates the TFs Lyl1 and Mnt which both induce Flt3 downregulation in aged LT-HSCs (Fig. 4 C), although the role of Flt3 in LT-HSCs, where it is lowly expressed and not present on the cell surface, is still relatively unexplored. Meanwhile, the 2^nd^ most highly ranked age-driving TF, Cebpd, is known to promote myeloid differentiation (Pawar et al., 2014) and both c-Maf1 and Cebpd are predicted to drive the upregulation of Gadd45g (Fig. 4C and Supplementary Fig. 4C), which has known roles in driving HSCs out of quiescence toward differentiation (Thalheimer et al., 2014). Furthermore, Atf3 (next in the ranked list) is known to prevent stem cell exhaustion (Liu et al., 2020), promote myeloid cell differentiation (Perrone et al., 2023) and both Cebpd and Atf3 are predicted in our differential GRN to directly downregulate the TF Bach1 (Supplementary Fig. 4A and B), which is known to suppress myelopoiesis (Itoh-Nakadai et al., 2014; Kato et al., 2018). Thus, inhibition of one or more of these TFs could potentially prevent the myeloid-biased output of aged murine LT-HSCs. It is also worth noting that the most highly ageing-upregulated TF is Rorb (which is also a LIG) and is predicted in the ageing-specific differential GRN to drive the downregulation of the TFs Dbp & Nr4a1 (Supplementary Fig. 4C), which have known roles associated with neuronal excitability, connectivity, and circadian rhythms in the brain (Kunst et al., 2015). Thus, our results predict a role for these neuron related TFs in LT-HSC ageing.

The TFs predicted to reprogram cells back to a youthful state in our analysis also have good evidence from the literature linking them to LT-HSC maintenance. For example, Gfi1 has roles in regulating differentiation and proliferation of LT-HSCs (Hock et al., 2004; Khandanpour et al., 2011), Hes1 is required to prevent stem cell exhaustion (Ma et al., 2020), while the role of Hif3a is less understood. In addition, Sox4 downregulation is thought to enable myeloid differentiation (Aue et al., 2011) and has known pioneer factor abilities (Katsuda et al., 2024). Our young LT-HSC-specific differential GRN predicts that Gfi1 & Hif3a both target Sox4 which upregulates the TF, Bach1 (Fig 5 D & E), which is known to supress myelopoiesis (Itoh-Nakadai et al., 2014; Kato et al., 2018). This appears to be the opposite role of Cebpd and Atf3, which are predicted to downregulate it, suggesting an antagonistic role of Gfi1 & Hif3a via Sox4 in combating age-associated gene expression changes driven by Cebpd and Atf3.

In summary, we have described epigenetic changes in aged murine LT-HSCs associated with the degradation of cell identity and expansion of myeloid-primed LT-HSCs. AP-1 factors and factors such as c-Maf, Cebpd and Atf3 are top age-driving factors, while Gfi1, Hif3a, Sox4 and Hes1 are strong candidates for restoring the youthful LT-HSC state. While the roles of these individual factors have been previously studied, often in the context of oncogenesis and blood cancers (Binod et al., 2024; Foronda et al., 2014; Hönes et al., 2017; Tian et al., 2013), TFs for cellular reprogramming are usually described as collections or cocktails of TFs due to their combined effect on the overall GRN, effects that are generally difficult to predict with single knockout or overexpression experiments. Future reprogramming experiments would likely benefit from assessing knockdown or overexpression of various combinations of our top ranked TFs to see if balanced and robust differentiation can be restored to aged murine LT-HSCs.

## Methods

### HSC, monocyte and NK cell isolation

For isolation of LT-HSCs, bone marrow from young (8-12 weeks old) and aged (100-130 weeks old) C57BL/6 female mice was harvested by extracting femur, tibia and pelvic bones, crushing with mortar and pestle in PBS with 2% FBS (FACS buffer) and filtering through 45 µm filter. Filtered bone marrow cells were then incubated with red cell removal buffer [RCRB; 156 mM NH4Cl (Sigma Aldrich), 11.9 mM, NaHCO3 (Sigma Aldrich) and 0.097 mM, EDTA, made in-house at WEHI] for 1 minute at room temperature (RT), pelleted at 1500 rpm, washed with FACS buffer, then lineage positive cells were depleted using the EasySep™ Mouse Hematopoietic Progenitor Isolation Kit (Stem Cell Technologies #19856) according to the manufacturer’s instructions. The bone marrow of approximately 12 young mice and 6 aged mice was pooled per experiment with 3 experimental (biological) replicates performed.

For isolation of monocytes and NK cells, spleens from young (8-13 weeks old) and aged (100-104 weeks) C57BL/6 female mice were harvested, placed in ice-cold PBS, mechanically dissociated using a 3 mL syringe plunger end, filtered through a 45 μm nylon mesh filter, incubated in 500 μL RCRB for 1 min at RT to remove red blood cells, then washed twice in ice-cold PBS. Centrifugation steps during red blood cell lysis and washing were performed at 1500 rpm at 4℃.

### LT-HSCs antibody staining and FACS

Lineage depleted cells were incubated with the following antibodies on ice for 30 mins: CD34-BV421 (Brilliant Violet 421™ anti-mouse CD34, BioLegend #152208) Flt3-APC (Ms CD135 APC A2F10.1, Becton Dickinson #560718) Sca1-BUV395 (Ms Ly-6A/E BUV395 D7 Becton Dickinson #563990) c-Kit-BV711 (Ms CD117 BV711 2B8 Becton Dickinson #563160) CD150-PE-Cy7 (PE/Cyanine7 anti-mouse CD150, TC15-12F12.2, BioLegend #115914) CD48-APC-Cy7 (APC/Cyanine7 anti-mouse CD150, HM48-1, BioLegend #103432) Lineage-biotin (Miltenyi Biotec Australia Pty. Ltd. #130-092-613) Cells were then washed and incubated with Streptavidin-BV650 (SAV BV650, Becton Dickinson #563855) on ice for a further 15 mins. For staining of mitochrondrial membrane potential (MMPot), the lineage depleted cells were then incubated with TMRM (Image-iT TMRM Reagent, ThermoFisher #I34361) and 50 mM Verapamil hydrochloride (99%, Merck, #V4629-1G) for 30 mins at 37°C before staining with SYTOX Blue (Thermo fisher #S34857) on ice for 20 mins. MMPot High LT-HSCs (CD34^-^ CD48^-^ Flt3^-^ CD150^+^ TMRM^High^ LSKs) and MMPot Low LT-HSCs (CD34^-^ CD48^-^ Flt3^-^ CD150^+^ TMRM^Low^ LSKs) were FACS sorted (Supplementary Fig. 1A, B and C), yielding approximately 400,000 aged and 60,000 young LT-HSCs per experiment.

### Monocyte & NK cell antibody staining and FACS

Splenocyte cell suspensions were incubated for 30 minutes on ice with the following antibodies: CD44-FITC (IM7.81, in-house, WEHI, Vic, Australia) B220-BUV395 (RA3-6B2, Becton Dickinson #563793) TCRβ-APC-eFluor 780 (H57-597, eBioscience #47-5961-82) Ly6C-eFluor 450 (HK1.4, eBioscience #48-5932-82) CD115-BV605 (T38-320, Becton Dickinson #743640) Ly6G-PE (1A8, Becton Dickinson #551461) CD4-APC (GK1.5-7, in-house, WEHI, Vic, Australia) NK1.1-PE-Cy7 (PK136, Becton Dickinson #552878) CD62L-biotin (MEL-14, Becton Dickinson # 553149) Cells were washed and stained for a further 15 mins with Streptavidin-Alexa594 (Thermo fisher #S32356), washed again and stained with Sytox blue (Thermo fisher #S34857) for 30 mins on ice. Monocytes (B220^-^ TCRβ^-^ CD115^+^ Ly6C^+^) and NK cells (B220^-^ TCRβ^-^ NK1.1^+^) were FACS sorted (Supplementary Fig. 1E) yielding approximately 320,000-540,000 aged and 200,000 young monocytes per experiment; and yielding approximately 360,000-640000 aged and 770,000 young NK cells per experiment with 2 experimental (biological) replicates in total.

### In situ Hi-C

Immediately following FACS, sorted MMPot High and Low LT-HSCs, or sorted monocytes and NK cells, were double fixed with 1% formaldhyde (10 mins) & 3mM disuccinimidyl glutarate (45 mins), pelleted at 850 × g at 4 °C, then snap frozen and stored at −80°C. Cells were then lysed in lysis buffer (10mM Tris-HCl pH8.0, 10mM NaCl, 0.2% Igepal CA630) with 50μl of protease inhibitors (Sigma, #P8340), washed with lysis buffer without protease inhibitors, then nuclei were placed in 0.5% sodium dodecyl sulfate (SDS) and incubated at 62°C for 10 minutes. The SDS was then quenched with 10% Triton X-100 (Sigma, #93443) for 15 mins at 37°C, then 100U of MboI restriction enzyme (NEB, #R0147) was added and the mixture incubated at 37°C overnight. The MboI enzyme was inactivated by heating to 62°C for 20 minutes, DNA fragment ends were biotinylated by incubating nuclei with biotin-14-ATP, dCTP, dGTP and dTTP and 5 U/μL DNA polymerase I Klenow (NEB, #M0210), then incubating at 37°C for 90 minutes with rotation. Proximity ligation was performed using 2000 U of T4 DNA Ligase (NEB, #M0202), incubating at room temperature for 4 hours with slow mixing on a rotator. Nuclei were lysed and crosslinked proteins were degraded by adding 50μl of 20mg/ml proteinase K (NEB, #P8107) and 120μl of 10% SDS, incubating at 55°C for 30 minutes, then 130μl of 5M sodium chloride was added and the mixture incubated at 68°C overnight with rotation.

### Hi-C DNA library preparation and sequencing

Hi-C DNA libraries were prepared by first purifying the biotinylated DNA: 1.6X volumes of pure ethanol and 0.1X volumes of 3M sodium acetate, pH 5.2, was added to each tube, mixed by inverting and incubated at −80°C for 15 minutes. DNA solutions were centrifuged at maximum speed for 15 minutes at 4°C, supernatant discarded, then precipitated DNA pellets washed twice with 70% ethanol, then resuspended in Tris buffer (10 mM Tris-HCl, pH 8) and incubated at 37°C for 15 minutes to fully dissolve the DNA. The purified biotinylated DNA was placed in 130μl Covaris microTUBEs (TrendBio, #520045) and sheared using the Covaris S220 (fill level: 10, duty cycle: 10, PIP: 175, cycles/burst: 200, time: 58 s). Sheared biotinylated DNA was purified using Dynabeads MyOne Streptavidin C1 beads (10mg/ml, ThermoFisher #65002) in Tween washing buffer (5mM Tris-HCl pH 7.5; 0.5mM EDTA; 1M NaCl; 0.05% Tween 20), DNA separated using a magnet, washed twice with buffer, then resuspended in 10 mM Tris-Cl pH 8.0. DNA libraries were end-repaired and Illumina sequencing barcodes added using the NEBNext Ultra DNA Library Prep Kit for Illumina (NEB, #E7645). The Hi-C libraries were then PCR amplified using the Phusion Taq polymerase (NEB, #M0530S) with P5+P7 primer cocktail (in-house, WEHI) and 9 cycles of PCR. Amplified DNA was then size-selected and purified using AMPure XP beads (Beckman Coulter, #A63881) and magnetic separation, then final libraries resuspended in 10mM TrisHCl pH 8. Libraries were prepared for 3 separate Hi-C experiments for LT-HSCs and 2 separate Hi-C experiments for monocytes and NK cells, then barcoded libraries were pooled. Pooled Hi-C libraries for LT-HSCs, or pooled monocytes and NK cells, were sequenced using the Illumina NextSeq 2000 sequencer on a P3 100 flow cell, using 66 bp paired-end reads.

### Hi-C analysis

Reads were aligned using Bowtie2 v.2.5.3 (Langmead & Salzberg, 2012), duplicates were marked using Picard tools v.3.3.0 (*Picard toolkit*, 2019), files sorted using Samtools v. 1.21 (Li et al., 2009). Differential analysis was performed on three replicates per condition for LT-HSC (combined low and high MMpot) and two replicates per condition for monocytes and NK cells, in two separate analysis using diffHic v.1.36.1 (A. T. Lun & G. K. Smyth, 2015). In brief, pairwise interactions were quantified at 100 kbp resolution using the squareCounts function. Genomic fragments were defined using an mm39 reference, and analyses were restricted to canonical chromosomes. Low-abundance and artifactual interactions were removed by applying filterDirect, which estimated non-specific ligation rates from inter-chromosomal pairs, and filterDiag to eliminate short-range (diagonal) contacts. For differential interaction analysis, systematic non-linear biases (including distance-dependent trends) were corrected by calculating offsets using the normOffsets function with method=“loess” from the csaw (Lun & Smyth, 2016) package. These offsets were calculated separately for near-diagonal (<500 kbp) and far-diagonal/inter-chromosomal interactions to account for distinct noise profiles. Differential interactions were then identified using the edgeR v.4.2.2 (Chen et al., 2025; Robinson et al., 2010) quasi-likelihood (QL) framework. Interaction data were converted into a DGEList, and biological variability was modelled through dispersion estimation with estimateDisp and glmQLFit. Differential interaction analysis between aged and young samples was then performed using glmQLFTest with subsequent multiple testing correction (FDR < 0.1), and significant interactions were annotated with genomic coordinates using GenomicRanges (Lawrence et al., 2013) and visualized using custom ggplot2 v.4.0.1 (Wickham, 2016) plots and using Integrative Genomics Viewer (IGV) v3.0.0 (Thorvaldsdóttir et al., 2013). Additional Hi-C heatmap and multitrack plots were made using the plotgardener v1.14.0 R package (Kramer et al., 2022), but for this the Hi-C data had blacklisted sites removed, replicate merged and counts normalized using Iterative Correction and Eigenvector decomposition (see the Methods section “Topologically associated domains (TAD) analysis” for details).

### Gene biological pathway enrichment analysis

Genes or DEGs that overlap regions of age-increased and age-decreased interactions were analysed using the R package clusterProfiler (Yu et al., 2012) by running the goenrich and enrichPathway functions with the default settings recommended in the documentation (p value 0.05, q value 0.2, with BH correction for the adjusted p value), and the results plotted using the dotplot and heatplot functions from the enrichplot v1.28.4 (Yu, 2025) package.

### Topologically associated domains (TAD) analysis

Hi-C data were processed using HiCExplorer v3.6 (Ramirez et al., 2018) for TAD analysis. Initially, mm39 aligned BAM files were coordinate-sorted using SAMtools v1.22.1 (Li et al., 2009) and reads overlapping mm39 blacklisted regions were removed via bedtools intersect. The filtered BAMs were then name-sorted and converted to pairs format using pairtools v1.1.3 (Open2C et al., 2024) with a minimum mapping quality of 30, a unique walks-policy, and a maximum inter-align gap of 20 bp. Subsequent steps included sorting, deduplication (allowing up to three mismatches), and filtering to retain valid intra- and inter-chromosomal interactions (with a minimum distance of 1 kb), thereby generating high-confidence interaction pairs. These pairs were converted into contact matrices in the .cool format, and replicates were merged using cooler merge. Finally, the merged .cool files were transformed into the HDF5 (.h5) format with hicConvertFormat.

The resulting Hi-C contact matrices were normalized using hicNormalize with the ‘smallest’ normalization method across multiple resolutions (ranging from 10 kb to 2 Mb) to ensure data comparability. Diagnostic plots were generated using hicCorrectMatrix, and matrices were further corrected using Iterative Correction and Eigenvector decomposition (ICE) normalization, with threshold values determined from the diagnostic plots. These ICE normalized, merged Hi-C files were for all Hi-C heatmap plots using the plotgardener v1.14.0 R package (Kramer et al., 2022). TADs were subsequently identified at 100kbp resolution using hicFindTADs, where the step size was set to the resolution and the minimum and maximum depths were defined as three- and ten-fold the resolution, respectively. TAD separation scores from hicFindTADs were then plotted for each condition (aged and young) and the significance of the difference in distribution determined using a Mann Whitney U test.

### Compartment analysis

Compartment analysis was performed using HiCDOC v.1.6.0 (Kurylo et al., 2023). Raw data stored in .cool files were first organized into HiCDOCDataSet objects by extracting resolution-specific replicates and conditions (aged vs. young) via the function HiCDOCDataSetFromCool. To ensure data quality, matrices were filtered using a series of functions: filterSmallChromosomes removed chromosomes below a set length threshold, filterSparseReplicates excluded replicates with fewer than 5% non-zero interactions, and filterWeakPositions eliminated bins with average interactions below 1/6th of the overall signal.

Normalization was performed in three sequential steps to correct for technical, biological, and distance-dependent biases. Technical biases were normalized using normalizeTechnicalBiases (employing cyclic loess normalization), followed by normalizeBiologicalBiases to adjust for sample-specific effects, and finally, normalizeDistanceEffect was applied with resolution-specific loess sample sizes to correct for the decay of interaction frequency with genomic distance. A/B compartments were then detected using detectCompartments and compartment differences between conditions were identified using the differences function (threshold = 0.05). For the final analyses, only the 200 kbp resolution data were used to generate compartment assignments, differential calls, and corresponding visualizations with plotCompartmentChanges.

### Shannon entropy analysis

Shannon entropy was calculated using a method similar to Jenkinson et al. (Jenkinson et al., 2017) by calculating the probability distribution of interactions of all 100 kbp bins within each 2 Mbp span. Entropy value distributions in young vs aged were then compared using a Mann Whitney U-test. In brief, MMPot high and low merged Hi-C BAM files (3 aged, 3 young) were initially processed using custom bash scripts to downsample the coordinate-sorted BAM files to a common depth: Unique read counts for each sample were determined and the minimum unique read count was used as a reference to calculate a downsampling fraction. BAM files were then subsampled using the SAMtools v1.22.1 (Li et al., 2009) view command with a fixed random seed (42) to ensure reproducibility, then a name-sorting step was performed to prepare the files for downstream analysis.

Downsampled Hi-C interaction matrices were imported into R as InteractionSet (Lun et al., 2016) objects. Bins were first filtered based on their size (e.g., retaining those within 90–110 kb) to maintain a consistent resolution. Genomic spans of 2 Mbp were then generated using the GenomicRanges tileGenome function (Lawrence et al., 2013), and bins were mapped to these spans via findOverlaps. Within each span, the interaction counts for each sample in each condition were aggregated by unique bin-pair identifiers so only aged and young samples were compared. Probabilities were computed from these counts, and Shannon entropy was calculated as the negative sum of the probability multiplied by the log₂ of the probability. This step was adapted from established protocols (Jenkinson et al., 2017; Liu et al., 2022) to capture the local interaction variability within defined genomic spans. Distribution of entropy values were compared between aged and young LT-HSCs using a non-parametric Mann Whitney U-test.

### RNA-seq analysis

RNA-seq data for aged and young LT-HSCs from Itokawa et al., 2022 (GSE162607) were analysed for differential expression as follows: Raw sequencing reads were subjected to quality control and adapter trimming. Adapters were removed and reads were filtered for a minimum length of 20 bp and a quality score of 20 using Cutadapt 4.9 (Martin, 2011). The quality of trimmed reads was further assessed using FastQC v0.11.x (Andrews, 2017). Cleaned reads were then aligned to the GRCm39 reference genome (Ensembl release 113) using STAR 2.7.11b (Dobin et al., 2013). Alignment was performed in two-pass mode, and output was sorted by coordinate into BAM format.

Gene-level read counts were quantified from the aligned BAM files using featureCounts from Subread v2.0.6 (Liao et al., 2013), with annotations provided by the _Mus musculus_ GRCm39.113 GTF file. Downstream differential expression analysis was conducted in R using the edgeR package 4.2.2 (Chen et al., 2025; Robinson et al., 2010). Gene annotation information, including Entrez Gene IDs and MGI symbols, was retrieved from Ensembl (mmusculus_gene_ensembl dataset) using the biomaRt R package (Durinck et al., 2009). A design matrix was constructed using the model.matrix function so that groups of the same age and cell type were compared (aged vs young LT-HSCs), then lowly expressed genes were filtered using filterByExpr, followed by Trimmed Mean of M-values (TMM) normalization to account for library size and compositional biases. A Quasi-Likelihood Negative Binomial Generalized Linear Model (GLM) was fitted, and differential expression was determined using Quasi-Likelihood F-tests. Genes with a false discovery rate (FDR) adjusted p-value of less than 0.05 were considered statistically significant. Genomic ranges for differentially expressed genes were subsequently incorporated using the rtracklayer v.1.68.0 R package (Lawrence et al., 2009), based on the same Ensembl GTF annotation. Importantly, LT-HSCs surface markers used for sorting cells in the GSE162607 dataset, i.e. Il-7^-^ CD34^-^ CD48^-^ Flt3^-^ CD150^+^ LSKs, are similar to those used for our Hi-C experiments, where Itokawa et al. use IL-7 to also isolate Il-7^-/+^ later progenitors CLPs, MEPs and GMPs (See (Itokawa et al., 2022)).

The differential expression values used to build gene regulatory networks were generated in R using DESeq2 v.1.48.2 (Love et al., 2014) on the STAR count matrices for LT-HSCs only. DESeqDataSet objects were constructed using DESeqDataSetFromMatrix, library size normalization was performed by estimating size factors with estimateSizeFactors and differential expression testing was then carried out using DESeq using alt_hypothesis = ‘greaterAbs’, independent_filtering = TRUE, p_adjust_method = ‘BH’ and minmu = 0.5. Pairwise contrasts were performed for aged/young or young/aged depending on whether the target condition was aged LT-HSCs or young LT-HSCs, respectively.

### Determination of lineage inappropriate expression

Raw RNA-seq expression counts for LT-HSCs, MPP1-4s, MEPs, GMPs and CLPs from the Itokawa et al. 2022 data (GSE162607) were normalized by a recently described technique called gene length corrected trimmed mean of M-values (GeTMM) (Smid et al., 2018), which uses edgeR’s TMM normalization on read counts that have been adjusted by gene length. By filtering for only genes that were significantly differentially expressed and setting upper and lower log2GeTMM expression thresholds we defined a set of genes lowly expressed or absent in young blood progenitors, which have low or negligible expression [log_2_(GeTMM) < 2.71] in young LT-HSCs, MPP1-4s, GMPs, MEPs or CLP and significantly higher expression [log_2_(GeTMM) > 2.88] in aged LT-HSCs. Restricting our definition again to genes that have near zero expression [log_2_(GeTMM) < 1] in young blood progenitors further defined genes we term “lineage inappropriate genes” (LIGs).

### ATAC-seq analysis

ATAC-seq FASTQ files from the Itokawa et al (2022) dataset (GSE162551) were quality-checked with FastQC v0.11.x (Andrews, 2017) and reads aligned to the GRCm39 mouse reference genome using Bowtie2 v2.5.3 (Langmead & Salzberg, 2012) and converted to BAM with SAMtools v1.22.1 (Li et al., 2009); alignments were coordinate-sorted and PCR duplicates were removed with Picard v3.1.1 (*Picard toolkit*, 2019) using MarkDuplicates (REMOVE_DUPLICATES=true). Deduplicated alignments were filtered to retain mapped reads with mapping quality ≥10, and mitochondrial reads were excluded; filtered BAMs were then indexed. To remove artefactual signal, reads overlapping genomic exclusion regions from mm39.excluderanges (Ogata et al., 2023) were removed with BEDTools v2.31.1 (Quinlan & Hall, 2010) and the remaining reads were re-sorted and indexed. Peaks were called per sample using MACS3 v3.0.1 (Zhang et al., 2008) with an effective genome size of 2,654,621,783 using the arguments --nomodel --shift −100 --extsize 200 with a significance threshold of q≤0.05.

Differential chromatin accessibility analysis was performed in R using DiffBind v3.18.0 (Stark & Brown, 2011) on blacklist-filtered ATAC-seq BAM files and MACS3 narrowPeak calls and analyses were restricted to primary autosomes (1–19) and chromosome X. A DiffBind sample sheet was constructed from the study metadata, and a consensus peakset was generated with minOverlap=2 to harmonize peak coordinates across samples. Read counts over consensus peaks were obtained with SummarizeOverlaps (bUseSummarizeOverlaps=TRUE) and, to reduce between-lineage variability, differential accessibility was performed separately for each hematopoietic cell type (LT-HSC, MPP1–4, GMP, MEP, CLP). For each cell type, an offset-based loess normalization was performed and differential testing was run as aged versus young using edgeR within DiffBind. Differentially accessible regions (DARs) were defined at FDR < 0.05.

### Motif enrichment analysis

Transcription factor (TF) binding motif enrichment analysis was performed on regions of differential interactions (DIs) identified in Hi-C data from aged versus young LT-HSCs at 100 kbp resolution. Briefly, DI regions were intersected with differentially accessible regions (DARs) determined from analysis of the Itokawa et al (2022) ATAC-seq dataset. Mouse TF binding motifs were retrieved from the HOCOMOCO v13 (Vorontsov et al., 2024) and database and filtered to only include TFs with a TPM > 0.1 in the Itokawa et al (2022) RNA-seq dataset, then converted to position weight matrices lists using the universalmotif v1.26.2 (Tremblay, 2024) and TFBSTools v1.46.0 (Tan & Lenhard, 2016) R packages. Enrichment of filtered TF motifs in DARs overlapping DIs was then performed using the monaLisa R package (Machlab et al., 2022). For enrichment of TF motifs in DARs overlapping DIs, only DARs and DIs that change in the same direction (age-opening DARs with gained DIs, or age-closing DARs with lost DIs) were analysed. To enable binning, a combined DAR-DI “logFC score” was calculated by multiplying together the log_2_FoldChange values for the DARs and the DIs they overlap, then making the age-closing/lost interaction scores negative by multiplying it by −1. The logFC scores were then binned into 4 bins and the DARs were analysed using the monaLisa calcBinnedMotifEnrR function. Results were filtered for motifs that were significantly enriched (enrichment adj.p < 0.05) then plotted using the plotMotifHeatmaps function.

### Network analysis and transcription factor ranking

In summary, TF-gene affinities were calculated using the STARE pipeline (Hecker et al., 2023) in gABC mode by combining Hi-C data and ATAC-seq data then mapping TF motifs to the ATAC-seq peaks and aggregating TF binding scores from different peaks assigned to the same gene. TF-gene affinities were then combined with RNA-seq expression values to generate GRNs. Differential GRNs were generated by subtracting the network weights of one condition from another and TFs were ranked, following the ANANSE pipeline (Xu et al., 2021), according to their strength of influence on the overall changes in network weights.

Firstly, Hi-C contact matrices that had previously been blacklist-filtered and ICE normalized were converted to STARE-compatible contact files. For each condition (young and aged LT-HSCs), Hi-C contact matrices at 100 kb in .h5 format were converted to a genome-wide GInteractions-style TSV using hicConvertFormat from HiCExplorer v3.6 (Ramirez et al., 2018) and then split by chromosome. For each chromosome, intra-chromosomal contacts were kept and written as a three-column STARE contact files (bin1, bin2, contact) at the chosen bin size (100 kb). ATAC-seq peak summits from all young and aged LT-HSC replicates were mereged, reduced to 1 bp summit positions, and expanded symmetrically by ±250 bp using the BEDTools v2.31.1 (Quinlan & Hall, 2010) slop command with mm39 chrom sizes to create a union set of fixed 500 bp windows. Windows overlapping mm39 blacklist regions were removed (bedtools subtract).

Per-peak activity was quantified by counting overlaps of each union window with filtered ATAC-seq BAM files using bedtools coverage - counts. The resulting peak-by-sample count matrix was imported into R and normalized with the csaw v1.42.0 (Aaron TL Lun & Gordon K Smyth, 2015) normOffsets function. Condition-specific chromatin accessibility was then summarized as offset normalized counts per million (CPM) for each replicate and as the mean CPM per condition, and written back to a BED-like table (chr, start, end, peak_id, per-replicate CPMs and Young/Old mean CPM).

These accessibility peak windows, together with the normalized Hi-C contacts and genome annotations, were then used as input to STARE in ABC mode to compute enhancer-gene ABC scores and TF-gene affinities. STARE was run via the provided STARE.sh driver script with the union ATAC-seq peak windows as -b, the Ensembl GRCm39 (mm39) primary assembly FASTA as -g, and the corresponding Ensembl GTF annotation as -a. For the ABC model we supplied the per-chromosome Hi-C contact files (-f), the Hi-C bin size (-k 100000), the column number for the mean ATAC-seq CPM value for each condition from per-peak activity table (-n), a gene-centred window of 5 Mb around TSSs (-w 5000000), an enhancer window of 5 Mb for the adapted ABC adjustment (-m 5000000), an ABC score cutoff of 0.02 (-t 0.02), and enabled both pseudocounts and the adapted ABC scoring (-d True, -q True) with tss_mode set to all_tss. Position-specific energy matrices (PSEMs) were generated from transfac-format PSCMs using the built-in PSCM-to-PSEM conversion. STARE was run using the HOCOMOCO-derived H13 CORE (Vorontsov et al., 2024) TF motif set. STARE first extracted open-chromatin sequences from the reference genome (bedtools getfasta), cleaned non-canonical bases, and then computed TF-DNA affinities for each peak using TRAP, which sums the per-position binding probabilities for each peak region, followed by integration of TRAP scores, ABC enhancer-gene scores and TSS annotations to produce TF-gene affinity matrices for each condition (young and old LT-HSCs). To make the STARE output compatible with ANANSE influence, the TF-gene affinity matrix values were scaled to a value of between 0 and 1. Mean TPM expression for target genes and TFs was then computed, rescaled to between 0 and 1, then these scaled expression values were used to modulate the scaled TF-gene affinity matrix scores (affinity_[0,1]_) such that each TF-gene edge weight becomes prob_TF,gene_ using the following:

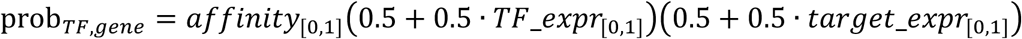

These expression-weighted affinity (prob) values were then used by the ANANSE influence command to rank the TFs in several steps using the DEG logFC values. First a differential network (Old - Young or Young - Old) was built by subtracting the source GRN from the target GRN (i.e. the modified STARE networks with the “prob” values as network edges), where only edges with positive differences (higher values in the target condition) were kept. For each TF, a Target score was then computed by summing, over all DEGs reachable within two steps in this differential network, the product of each TF’s target gene connection weights and the target gene expression-change score (i.e. the |log₂FC|):

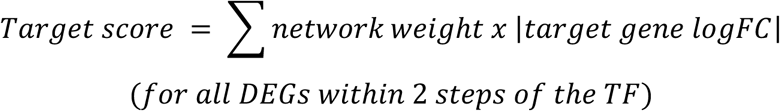

A G_score equal to the TF’s own expression-change score (|log₂FC| if upregulated and padj < 0.05) was also assigned:

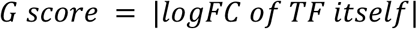

Finally, ANANSE influence ranked TFs by an Influence score, whereby the rank is equivalent to:

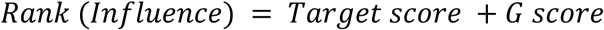

The TF Rank (Influence score) is derived by summing the Target score and G score then scaling to a value between 0 and 1. A TF is thus highly ranked if it is strongly connected to multiple target DEGs (1 or 2 steps away in the network) that have high changes in expression and if the TF itself is strongly upregulated, but whereby the connectivity to multiple DEGs generally has more influence in the final ranking.

Differential GRN plots were generated in R by importing the TF influence table and differential network (source, target, weight) with readr v 2.1.5 (Wickham et al., 2024) and dplyr v 1.1.4 (Wickham et al., 2023), constructing directed graphs with igraph v 2.2.1 (Csardi & Nepusz, 2006), then visualising them with ggraph v 2.2.2 (Pedersen, 2025) and ggplot2 v4.0.1 (Villanueva & Chen, 2019). For each TF of interest, a TF-centered network was extracted to a maximum path length of two steps (max_steps = 2) to reflect ANANSE’s influence neighbourhood, and edges were restricted to outward connections only to emphasise TF→TF→gene cascades. To limit visual complexity, the differential network was filtered to the strongest edges (weight ≥ 1×10⁻⁴) and plotted subnetworks retained TF nodes plus differentially expressed target genes defined from DESeq2 results (padj ≤ 0.05 and |log2FoldChange| ≥ 0.2). The networks weights are shown in Supplementary Table 4.

## Supplementary Figures

**Supplementary figure 1.**
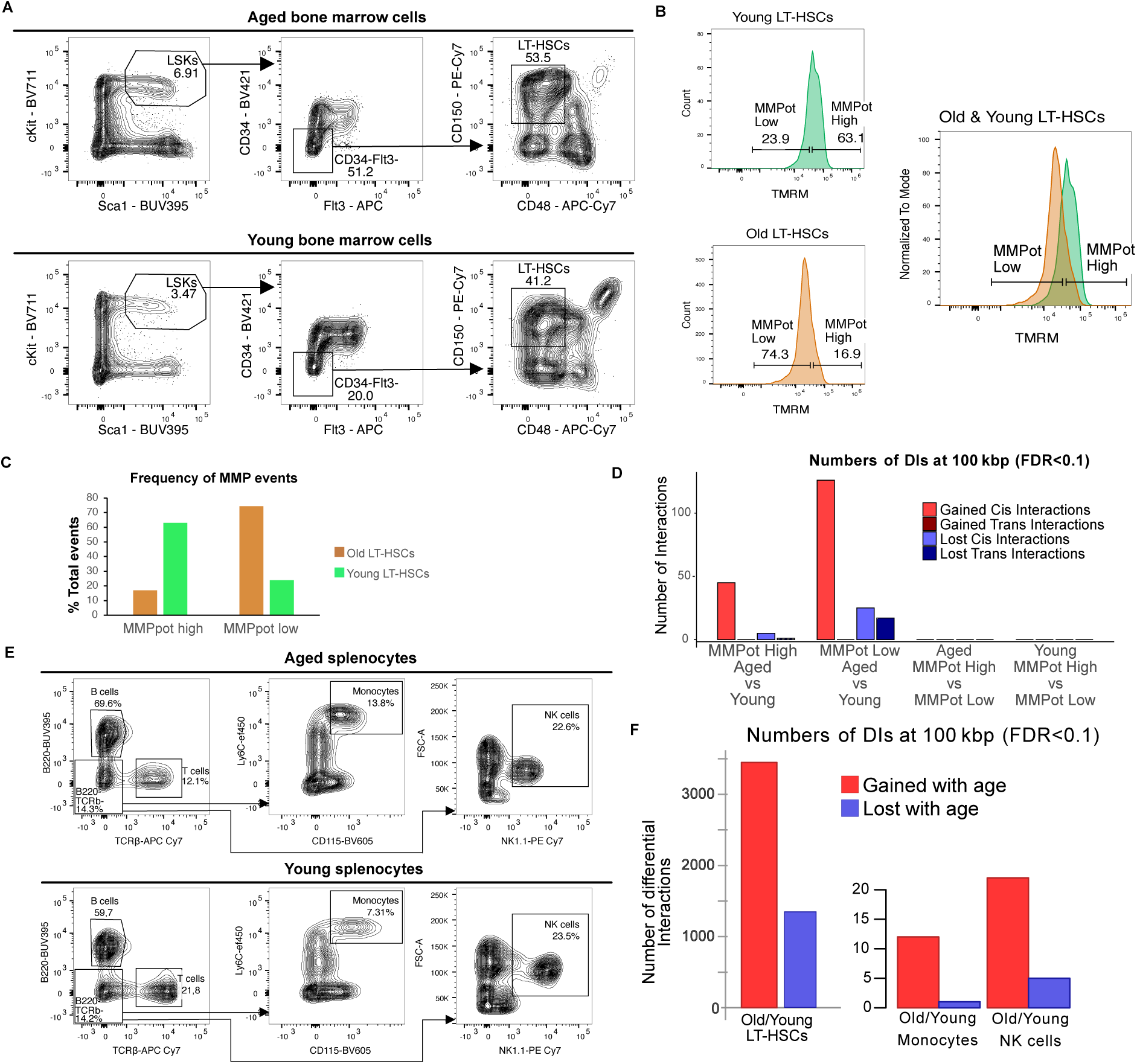
Sorting strategy and differential interaction results for murine (C57Bl/6) aged and young mitochondrial membrane potential (MMPot) high and low long-term hematopoietic stem cells (LT-HSCs), monocytes and NK cells. A. Fluoresence associtated cell sorting (FACS) plots for the isolation of aged and young LT-HSCs; B. FACS plots for isolation of aged (old) and young MMPot high and low LT-HSCs; C. Bar plot of the typical frequency (%) of MMPot high and MMPot low LT-HSCs obtained from each age condition; D. Differential interaction (DI) numbers for the four different contrasts tested, comparing either age or MMPot status, showing that ageing has a stronger effect than MMPot status; E. FACS plots for the isolation of murine (C57Bl/6) aged and young monocytes and NK cells; F. DI numbers for aggregated aged (old) versus young MMPot combined LT-HSCs (left) compared to old versus young monocytes or NK cells, showing that these mature cell types acquire far less chromatin architecteral differences than LT-HSCs during ageing.

**Supplementary figure 2.**
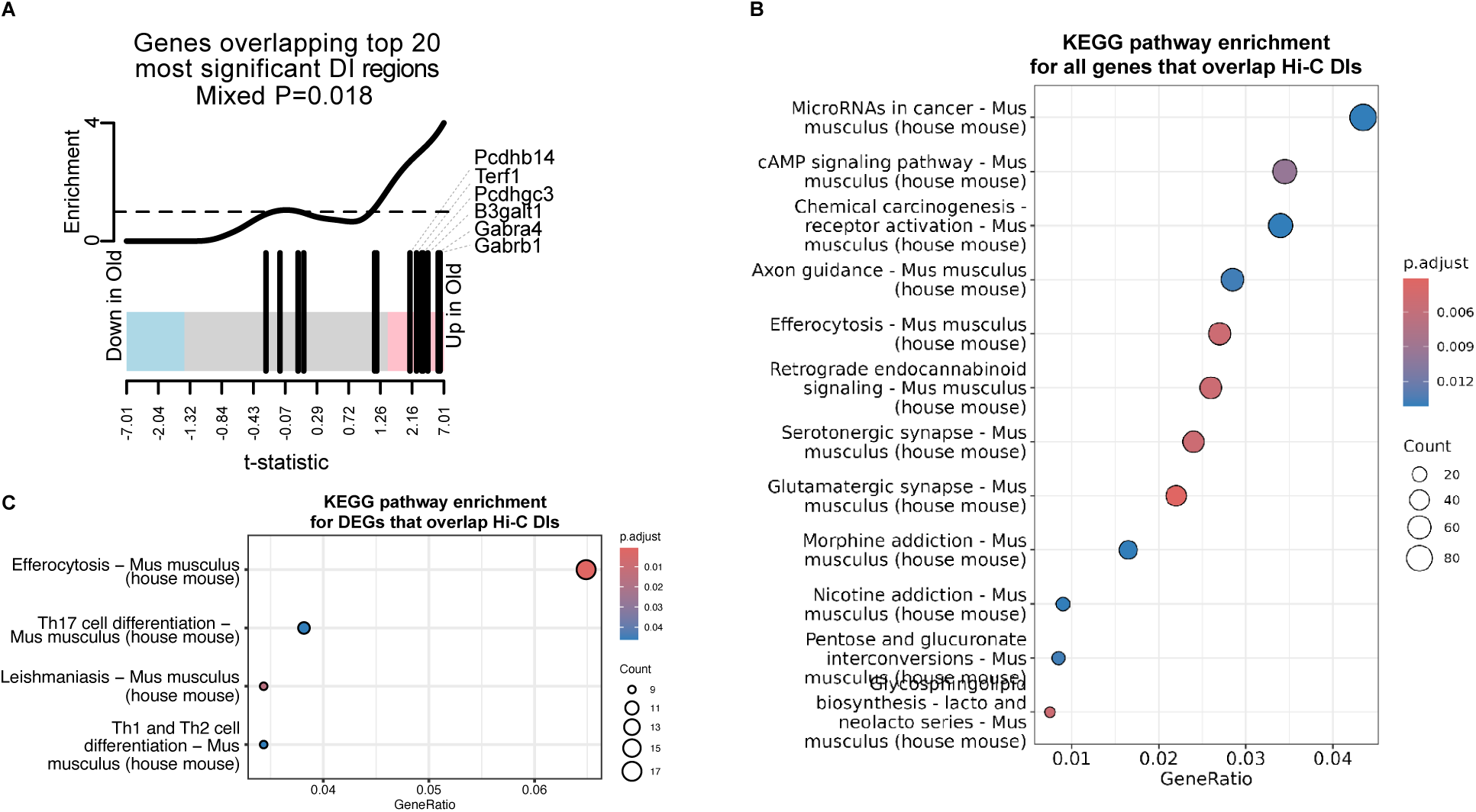
Fry test and KEGG pathway enrichment analysis results. A. Fry test showing that differentially expressed genes (DEGs) are enriched in the top 20 differentially interacting (DI) regions that alter interaction frequency with age and that the upregulated DEGs B3galt1, Gabra4, Gabra1 and Terf1 are particularly enriched in these top DI regions; B. Kyoto Encyclopedia of Genes and Genomes (KEGG) pathway enrichment analysis results for all genes that overlap HI-C DI regions that change with age in long-term hematopoietic stem cells (LT-HSCs); C. KEGG pathway enrichment analysis results for all DEGs that overlap HI-C DI regions that change with age in long-term hematopoietic stem cells (LT-HSCs).

**Supplementary figure 3.**
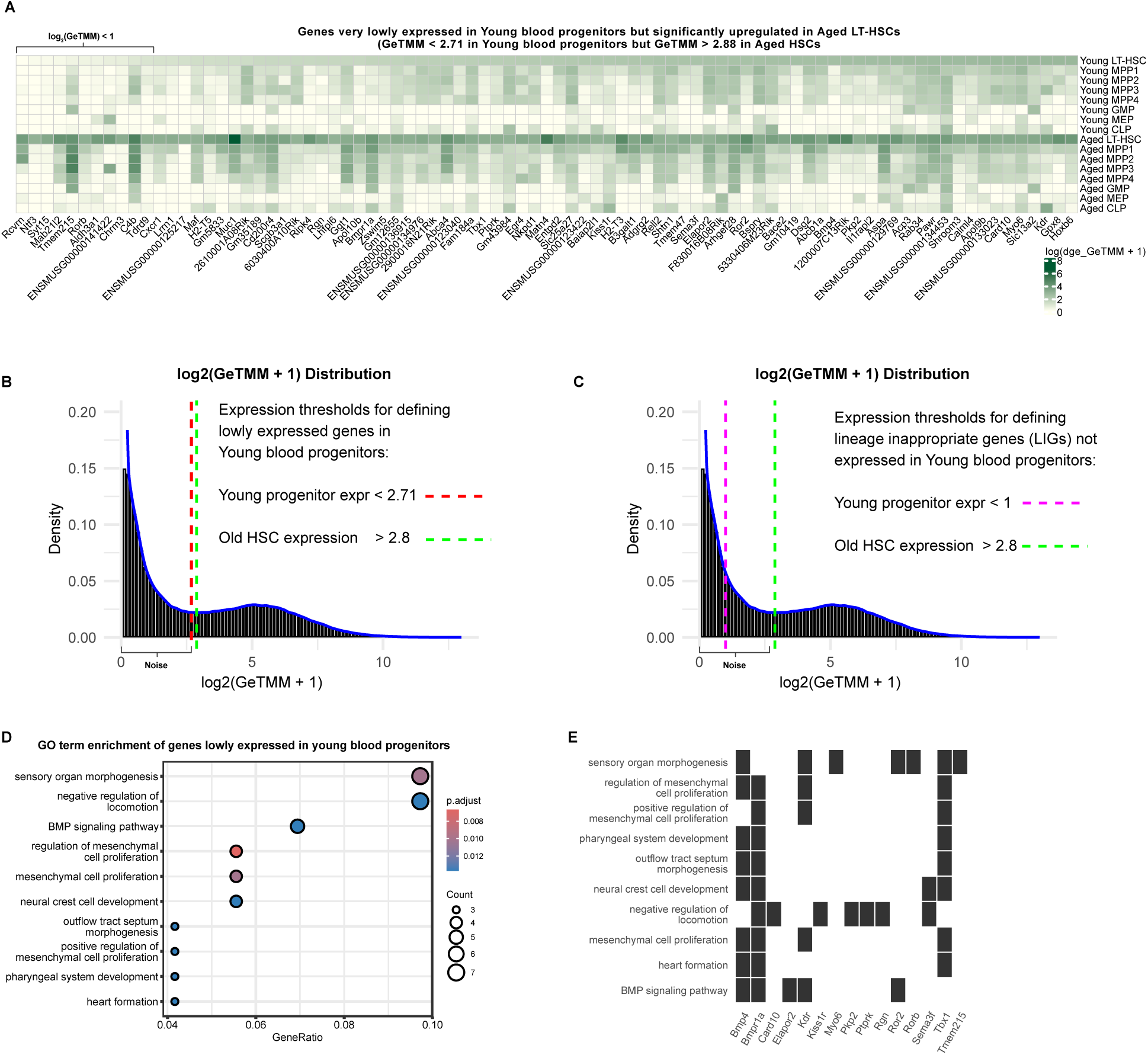
Significantly upregulated genes in aged long-term hematopoietic stem cells (LT-HSCs) that are absent or lowly expressed in young blood progenitors. A. Heatmap of log_2_ transformed gene-length corrected trimmed mean of M-values (GeTMM) expression values for 85 genes that appear to be lowly expressed or not-expressed in young blood progenitors (GeTMM expression value < 2.71) but that are significantly upregulated in aged LT-HSCs (GeTMM expression value > 2.88); B. Distribution of log_2_ transformed GeTMM expression values to determine express value thresholds that define actual expression versus noise where genes defined as “expressed” were above the 2.88 threshold (green dashed line) and genes defined as “not expressed” or “very lowly expressed” were below the 2.71 threshold (red dashed line); C. Distribution of log_2_ transformed GeTMM expression thresholds for lineage inapproriate genes (LIGs) whereby the upper threshold of 2.88 remained the same (green dashed line) but the lower threshold for the “not expressed” genes was lowered to the log_2_ GeTMM value of 1 (pink dashed line); D. Gene ontology (GO) term enrichment results for the genes shown in panel A; E. GO terms and the gene sets that that are enriched in each term.

**Supplementary figure 4.**
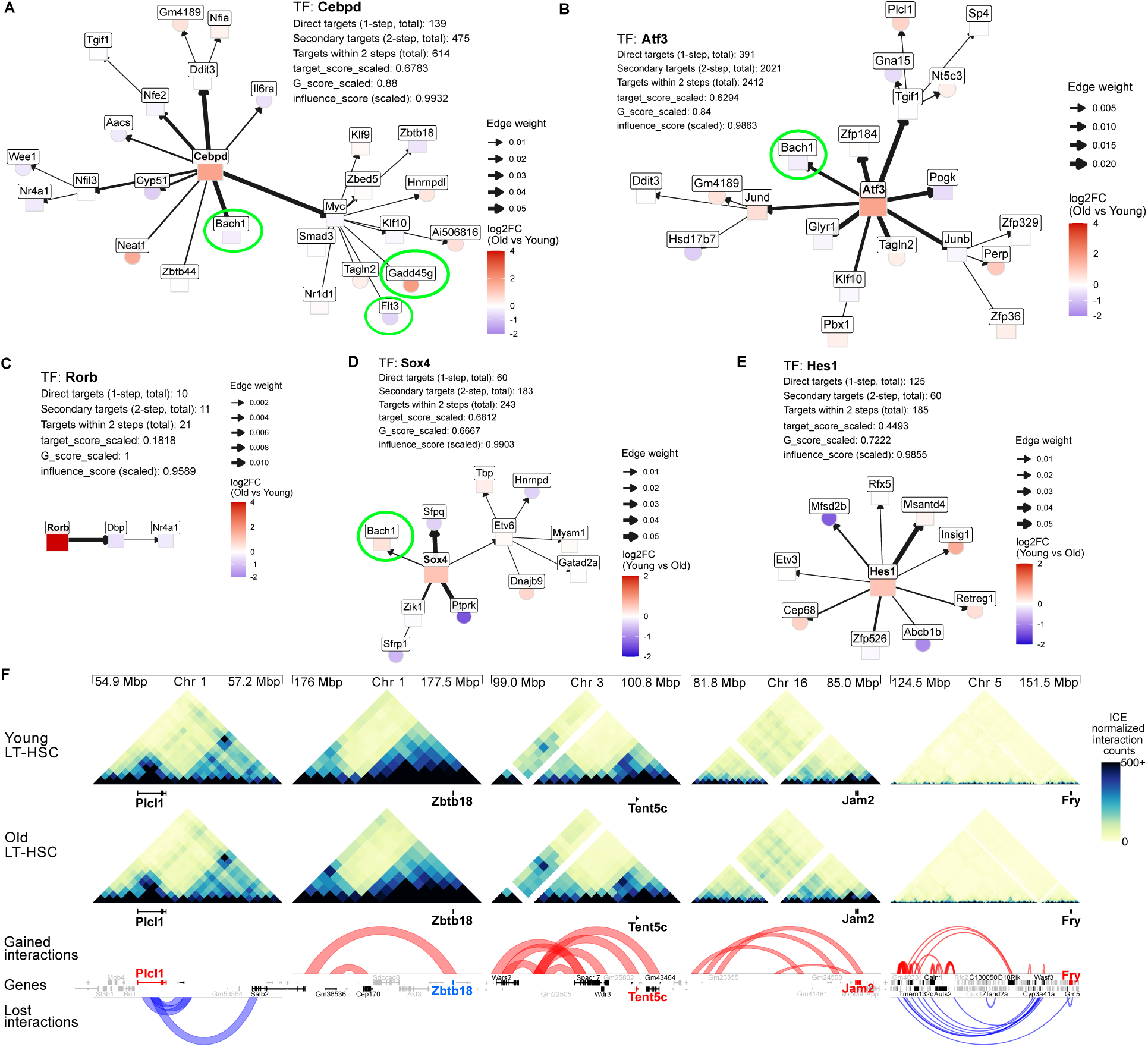
A. Simplified network diagram of the 2nd ranked age-driving TF, Cebpd, which is predicted to regulate 139 direct targets in this differential gene regulatory network (GRN), including the downregulation of the myeloid supressing TF Bach1 and regulation of Myc which in turn regulates Flt3 downregulation and Gadd45g upregulation (shown in green circles); B. Simplified network diagram of the 3rd ranked age-driving TF, Atf3, which is predicted to regulate 391 direct targets in this differential GRN, including the downregulation of Bach1 (shown in green circles) and upregulation of AP-1 factors such as Jund; C. Simplified network diagram of the 7th ranked age-driving TF, Rorb, which is predicted to regulate 10 direct targets in this differential GRN, including the downregulation of the TFs Dbp and Nr4a1; D. Simplified network diagram of the 3rd ranked youth-promoting TF, Sox4, which is predicted to regulate 60 direct targets in the young vs aged differential GRN, including the upregulation of the myeloid supressing TF Bach1 (shown in green circle); E. Simplified network diagram of the 4rd ranked youth-promoting TF, Hes1, which is predicted to regulate 125 direct targets in the young vs aged differential GRN, including the downregulation of the sphingosine-1-phosphate transporter gene, *Mfsd2b*, and upregulation of the suppressor of cholesterol synthesis gene, *Insig1*; F. Hi-C heatmaps and arc-plots for Young and Aged LT-HSCs showing region interactivity that changes with age near the differentially expressed genes (DEGs) targeted by highest ranked age-driving TF, c-Maf, including the DEGs *Plcl1*, *Zbtb18*, *Tent5c*, *Jam2* and *Fry* (DEGs in red are upregulated, DEGs in blue are downregulated).

